# Mammalian Pumilio Proteins Control Cellular Morphology, Migration, and Adhesion

**DOI:** 10.1101/2022.07.19.500580

**Authors:** Erin L. Sternburg, Jordan J. Lillibridge, Rattapol Phandthong, Fedor V. Karginov

## Abstract

Pumilio proteins are RNA-binding proteins that control mRNA translation and stability by binding to the 3’ UTR of target mRNAs. Mammals have two canonical Pumilio proteins, PUM1 and PUM2, which are known to act in many biological processes, including embryonic development, neurogenesis, cell cycle regulation and genomic stability. Here, we characterized a new role of both PUM1 and PUM2 in regulating cell morphology, migration, and adhesion in T-REx-293 cells, in addition to previously known defects in growth rate. Gene ontology analysis of differentially expressed genes in PDKO cells for both cellular component and biological process showed enrichment in categories related to adhesion and migration. PDKO cells had a collective cell migration rate significantly lower than that of WT cells and displayed changes in actin morphology. In addition, during growth, PDKO cells aggregated into clusters (clumps) due to an inability to escape cell-cell contacts. Addition of extracellular matrix (Matrigel) alleviated the clumping phenotype. Collagen IV (ColIV), a major component of Matrigel, was shown to be the driving force in allowing PDKO cells to monolayer appropriately, however, ColIV protein levels remained unperturbed in PDKO cells. This study characterizes a novel cellular phenotype associated with cellular morphology, migration, and adhesion which can aid in developing better models for PUM function in both developmental processes and disease.

## Introduction

Post-transcriptional regulation is fundamental to proper control of protein expression. This control is executed by trans-acting RNA binding factors, including RNA binding proteins (RBPs). Pumilio proteins belong to a broad group of RBPs that control mRNA translation and stability by binding to the 3’ UTR. As members of the highly conserved PUF family, Pumilio proteins have been studied in organisms from yeast to humans (1). Pumilio was originally characterized in Drosophila as a key regulator in embryonic development (2-4), where it acts cooperatively with nanos to regulate protein expression of the embryonic patterning gene, hunchback. Since its discovery, Pumilio proteins have been extensively characterized in invertebrates, including flies, worms, and yeast. These studies have highlighted the evolutionarily conserved function of Pumilio proteins in regulating development and germline maintenance (5-8).

Mammals have two canonical Pumilio proteins, PUM1 and PUM2, which are highly conserved across organisms. The two mammalian homologs share 69% identity / 74% similarity along their entire length. At the C-terminus is the ∼340 amino acid Pumilio homology domain (Pum-HD) that is responsible for sequence-specific RNA binding. Most of the differences among Pumilio homologs in different organisms, as well as between the two human paralogs, lie within the N-terminal region. PUM1 contains an extended N-terminal region, which has been hypothesized to recruit other binding partners and to be the site of additional regulation. Similar to many RBPs, significant N-terminal portions of both PUMs are predicted to be disordered and likely play a role in recruitment to phase-separated droplets under various cellular conditions (9). The two paralogs are largely redundant, although differences in target binding do exist (10-12). Both proteins recognize target transcripts by binding to a conserved Pumilio recognition element (PRE) 5′-UGUAnAUA-3′ (4, 13, 14). When bound, Pumilio proteins repress target transcripts through the recruitment of machinery which can inhibit translation and destabilize the transcript (11, 15). Similar to Drosophila, NANOS paralogues are thought to play a role in modulating the binding and regulatory behavior of PUMs (16, 17).

In recent years, the role of Pumilio proteins in mammals has been studied, validating many conserved PUM functions and identifying new ones. PUM proteins are essential to mammalian embryonic development; double knockout mouse embryos are inviable due to an inability to complete gastrulation (18, 19). PUM is also present in mouse oocytes and is demonstrated to have a maternal effect during early embryogenesis (20). Pumilio proteins also play an important role in gametogenesis (21, 22). For both oogenesis and spermatogenesis, Pumilio proteins regulate transcripts that control self-renewal of early germ cell populations.

Pumilio proteins also play a substantial role in neurogenesis and in proper neuronal function. Brain-specific PUM double knockout mice show defects in brain development (19) and PUM2 has been demonstrated to help specify cell fate in neuronal stem cells (23). Loss of PUM1 has been shown to increase the expression of Ataxin1, a defect that, in both mice and humans, causes spinocerebellar ataxia (24, 25). Within the immune system, Pumilio proteins regulate genes involved in innate immunity (26) and in the proper maintenance of hematopoietic stem cells (27). In addition, PUM proteins are known to regulate transcripts of proteins necessary for cell cycle regulation and genome stability (28-31).

Cells control their morphology, migration, and adhesion to each other and to the extracellular matrix (ECM) through interconnected regulatory pathways. Proper ECM deposition and sensing is imperative for cells to respond effectively to their microenvironment, and the ECM plays important roles in cell-cell signaling, motility, proliferation, and differentiation (32-34). The cellular-extracellular matrix interface has been rigorously studied for decades, and many context-dependent mechanistic relationships have been elucidated. For example, when epithelial tissue interactions with its cognate ECM are altered, reduced adhesion and misregulated morphology can lead to malignant behavior (35). In a neuronal cell line, an ECM component is a necessary signal for cell-cycle exit and neurite formation during differentiation, affecting the cells’ overall morphology (36). The function of RBPs in regulating these cellular processes are not well understood. Here, we demonstrate a role of mammalian PUM proteins in regulating cell morphology, migration, and adhesion though characterization of PUM double knockout (PDKO) T-REx-293 cells. We validate that the observed phenotype is PUM-dependent and provide further evidence that PUM proteins regulate transcripts important in cell adhesion and migration pathways.

## Results

### Double knockout of PUM1 and PUM2 affects the growth rate of T-REx-293 cells

PUM double knockout T-REx-293 cells were previously generated (10) and shown to lack PUM1 and PUM2 protein expression (Figure 1A). The PDKO cells displayed a diminished growth rate compared to WT (Figure 1B). This growth phenotype was also observed in HCT116 PDKO cells (30) (Supplementary Figure 1A).To confirm that the effect on growth rate is PUM-dependent, PUM1 and PUM2 were individually reintroduced into T-REx-293 PDKO cells (along with a GFP control), stable integrant cell populations were selected, and PUM expression was confirmed by western blot (Figure 1A). PUM1 rescue increased growth to a level that is significantly higher than that of PDKO (Figure 1C). PUM2 showed a similar, but not statistically significant, increase. However, rescue by PUM1 or PUM2 individually did not fully restore the growth rate to WT levels. Dosage effects of total PUM protein have been reported to affect neurodevelopment and genomic stability (24, 25, 30), and could explain the partial rescue. This observation is also in agreement with the notion that PUM proteins have non-redundant functions, with both proteins being necessary for WT function to be restored. Thus, PUM1+2 deletion in different cellular contexts leads to a decreased growth rate, and can be rescued by PUM1.

**Figure 1:**
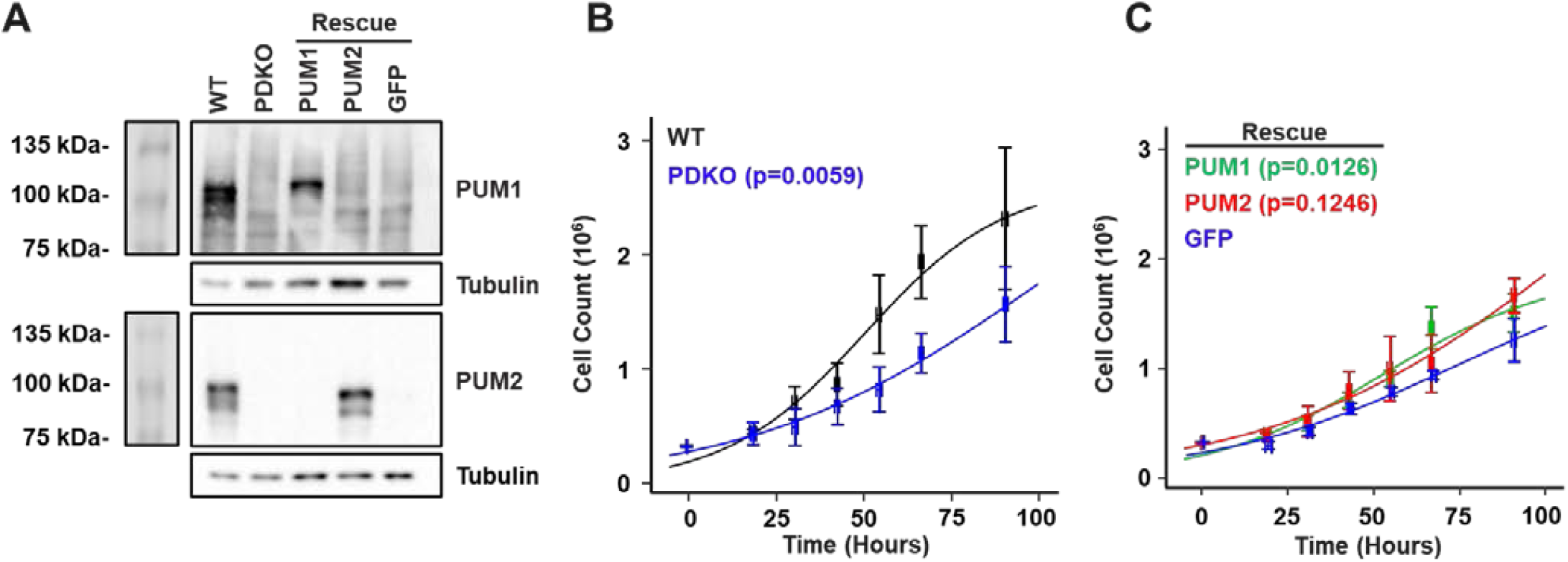
Cell growth rate is modulated by both PUM proteins. (A) Western blot for PUM1 and PUM2 in WT, PDKO, and PDKO cells stably transfected with PUM1, PUM2, or GFP. Tubulin serves as a control. (B) Growth rates of WT (black) and PDKO (blue) T-REx-293 cells. (C) Growth rates of PDKO T-REx-293 cells stably transfected with PUM1 (green), PUM2 (red), or GFP (blue). Growth measurements were calculated from three biological replicates, with error bars representing standard error of the mean. Logistic growth model fits of the data (trend lines) were compared using ANOVA F-test to determine statistical significance.

### PUM proteins extensively regulate mRNA levels and control cell movement and adhesion processes

To examine the genes and pathways that are regulated by PUM proteins, RNA-sequencing was performed on the above T-REx-293 and HCT116 cell lines. In T-REx-293 cells, gene expression in the WT line was the most distinct from the rest (Supplementary Figure 2A), and both PDKO and GFP+PDKO (negative rescue control) lines showed > 1000 differentially expressed (DE) genes compared to WT, while not being substantially different from each other (Supplementary Figure 2B). Thus, the PDKO and GFP-PDKO data were considered together (denoted PDKO in this section) in their comparisons to WT and PUM1/2 rescue lines. Deletion of PUM1+2 led to differential expression of 1706 genes (at a 2-fold and <5% FDR cutoff) compared to WT (Figure 2A). Among these, a significant majority (1188) were upregulated and 518 were downregulated, consistent with the predominantly repressive role of PUM proteins. Knockout of PUM1+2 in HCT116 cells revealed 485 DE genes, again predominantly upregulated (Supplementary Figure 2B).

**Figure 2:**
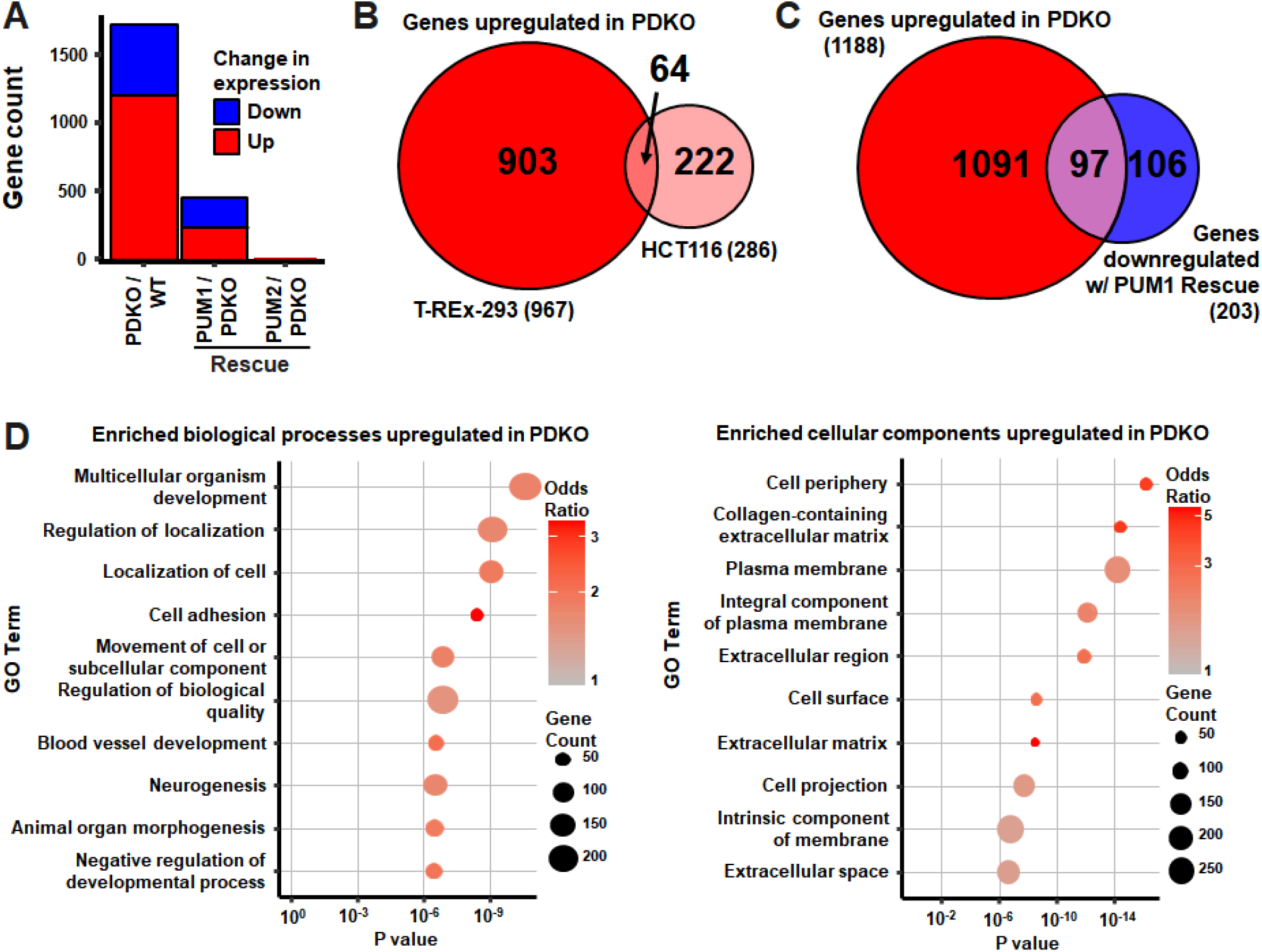
RNAseq analysis indicates that. PUM proteins extensively regulate mRNA levels and control cell movement and adhesion processes. (A) Number of differentially expressed genes in T-REx-293 PDKO vs WT, and PUM1 or PUM2 rescue vs PDKO comparisons. (B) Overlap among genes upregulated in T-REx-293 and HCT116 PDKO cells. (C) Overlap of genes upregulated in T-REx-293 PDKO vs WT with genes downregulated in PUM1 rescue vs PDKO. (D) Gene ontology (GO) biological process and cellular component terms enriched in genes upregulated in PDKO vs WT cells.

To determine whether PUM disruption regulates similar sets of genes in different cell types and settings, we compared our DE gene sets with each other and with previously published data. A strongly significant overlap in upregulated genes was identified between the T-REx-293 and HCT116 cells (Fisher’s exact test p-value 1.5 × 10^−27^, odds ratio 6.6; Figure 2B), while a weaker overlap was observed among the downregulated genes (p-value 0.03, odds ratio 2.4; Supplementary Figure 2C). Similarly, comparison with PUM1+2 knockdown in 293 cells (11) (at the same statistical cutoffs) revealed a robust overlap in upregulated genes (p-value 2.9 × 10^−11^, odds ratio 13.6; Supplementary Figure 2D), with some overlap for downregulated genes (p-value 0.001, odds ratio 10.5; Supplementary Figure 2E). Thus, PUM proteins exert their repressive functions on a consistent subset of genes across the examined datasets. The mRNAs that are stabilized by PUM proteins show more variability between cell types, potentially due to indirect effects or contributions from cell-type specific regulatory factors.

Importantly, rescue of the T-REx-293 PDKO line with PUM1 or PUM2 induced changes in gene expression. The impact of PUM1 was substantially broader than that of PUM2, with 436 and 10 DE genes, respectively (Figure 2A). Interestingly, nearly half of the genes downregulated in PUM1 rescue cells (compared to PDKO) were initially upregulated in PDKO relative to WT (p-value 3.6 × 10^−75^, odds ratio 21.0; Figure 2C). Conversely, a substantial portion of genes downregulated in PDKO vs WT were upregulated upon PUM1 rescue (p-value 4.3 × 10^−37^, odds ratio 15.0; Supplementary Figure 2F). These reciprocal changes indicate that reintroduction of PUM1 into the double knockout substantially reverts the gene expression program toward the WT state. Nevertheless, the majority of DE genes between PDKO and WT cells were not restored, suggesting additional mRNA-specific targeting by PUM2.

To identify the cellular functions and processes affected by PUM proteins, we performed gene ontology (GO) analysis, identifying a large number of enriched gene categories. For example, genes upregulated in PDKO were associated with 129/42/27 biological function (BP), cellular component (CC), and molecular function (MF) categories, respectively. The top 10 BP and CC categories are shown in Figure 2D. Interestingly, PUM proteins impacted cell adhesion and movement processes, and PUM-regulated proteins were enriched in those localized to the cell periphery/surface/membrane, as well as the ECM, including collagen-associated ECM. Together, these results strongly suggested that PUM proteins control aspects of cell movement, adhesion and/or interaction with the ECM.

### PUM proteins regulate actin morphology and alter cell migration rate

To investigate potential roles of PUM in adhesion and migration, we examined whether any structural changes were associated with the cytoskeleton of PDKO cells. To this end, WT and PDKO T-REx-293 cells were transfected with a Utrophin-RFP marker to label filamentous actin structures. Fluorescence images of sparsely growing live cells were blindly scored based on their major actin cytoskeletal structure related to locomotion: lamellipodia (flat, usually broad, plate-like extensions), filopodia (long, slender extensions), or other (does not fit either other description). Representative images are shown in Figure 3A. WT cells displayed a strong predominance of filopodia structures (Figure 3B). In contrast, PDKO cells showed a significantly different distribution with an even amount of filopodia- and lamellipodia-dominant cells. These results indicate that PUM proteins impact the cytoskeletal behavior of T-REx-293 cells, which could underlie potential defects in motility.

**Figure 3:**
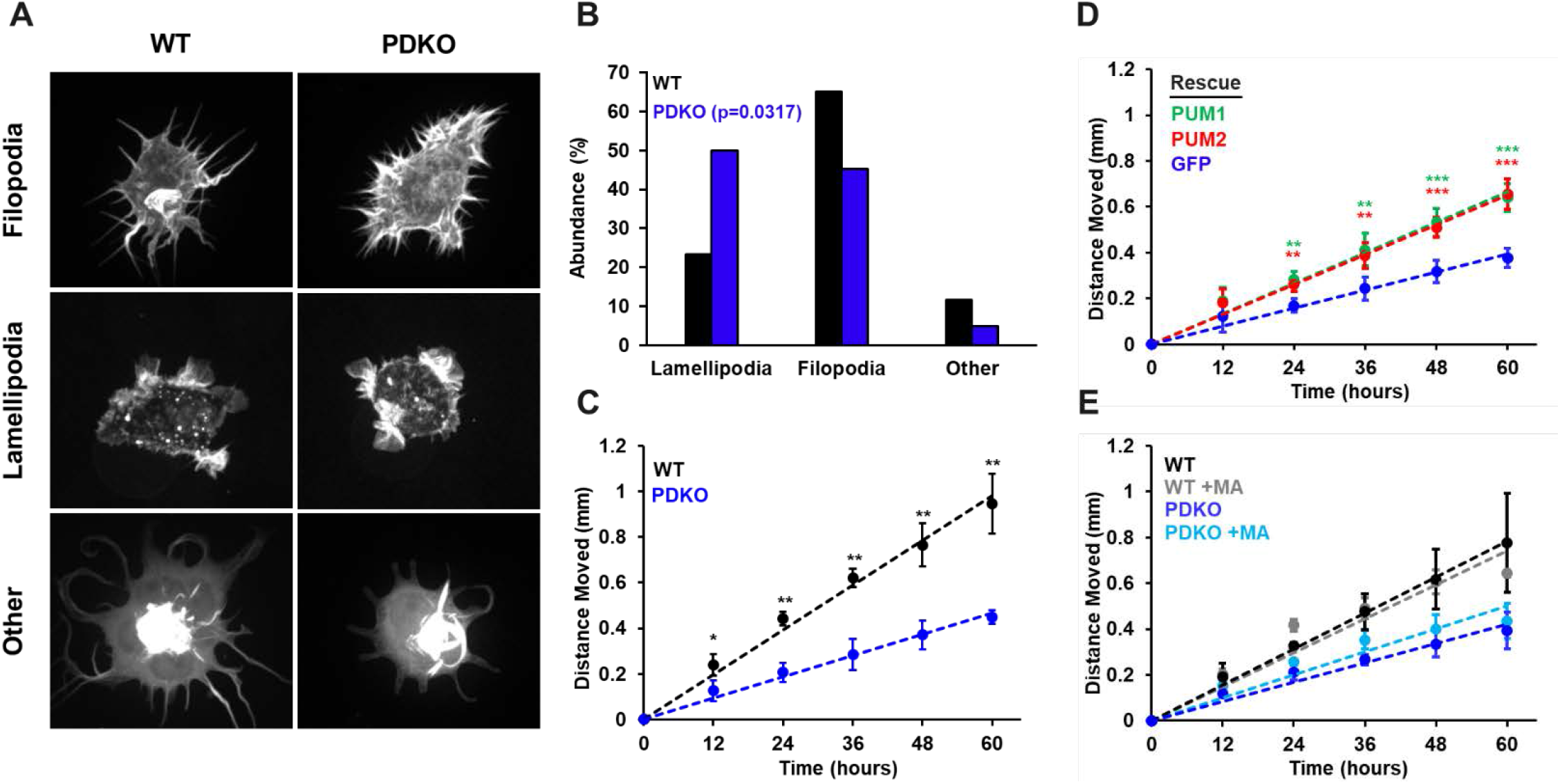
PUM1 and PUM2 regulate actin morphology and alter migration rate. (A) Representative images of WT and PDKO T-REx-293 cells transfected with mRFP-UtrCH to label filamentous actin. (B) Percent abundance of cells categorized to primarily contain lamellipodia, filopodia, or other structures. Cell images were scored blindly. Distributions were compared using a Chi-squared test. (C) Clonal ring assay migration rates of WT and PDKO cells. (D) Migration rates of PDKO cells rescued with PUM1, PUM2, or GFP. (E) Clonal ring migration rates of WT and PDKO cells with and without Matrigel. Matrigel was plated at a 1/50 dilution. Measurements were calculated over 3-5 biological replicates. Statistical significance was determined by Student’s T-test. *= p<0.05, **= p<0.01, ***= p<0.001.

To investigate the influence of PUM proteins on cell migration, we performed a clonal ring migration assay. Cells were plated inside a cloning ring and given time to adhere at full confluency. The cloning ring was then removed, and collective cell movement into the surrounding free space was measured. PDKO (and GFP control) cells migrated at a significantly slower rate than that of WT cells (Figure 3C), and migration of cells was partially restored in PUM1 or PUM2 rescue cell lines (Figure 3D). Interestingly, HCT116 cells also showed growth rate defects upon loss of PUM, but did not show any difference in migration rate between WT and PDKO cells (Supplementary Figure 1A and 1B). Taken together, PUM1 and PUM2 affect cytoskeletal cell morphology and migration in T-REx-293 cells, and this defect is separable from the growth rate defect.

### PUM1 and PUM2 affect cell-cell adhesion and separation

During regular culture, PDKO cells were observed to aggregate and form clusters after 2-3 days, in contrast to the even monolayer typical of WT T-REx-293 cells (Figure 4A). The effect could not be attributed to the slowed growth rate of PDKO cells, since even when cultured for extended periods of time, PDKO cells were unable to completely fill in the surface of a plate.

**Figure 4:**
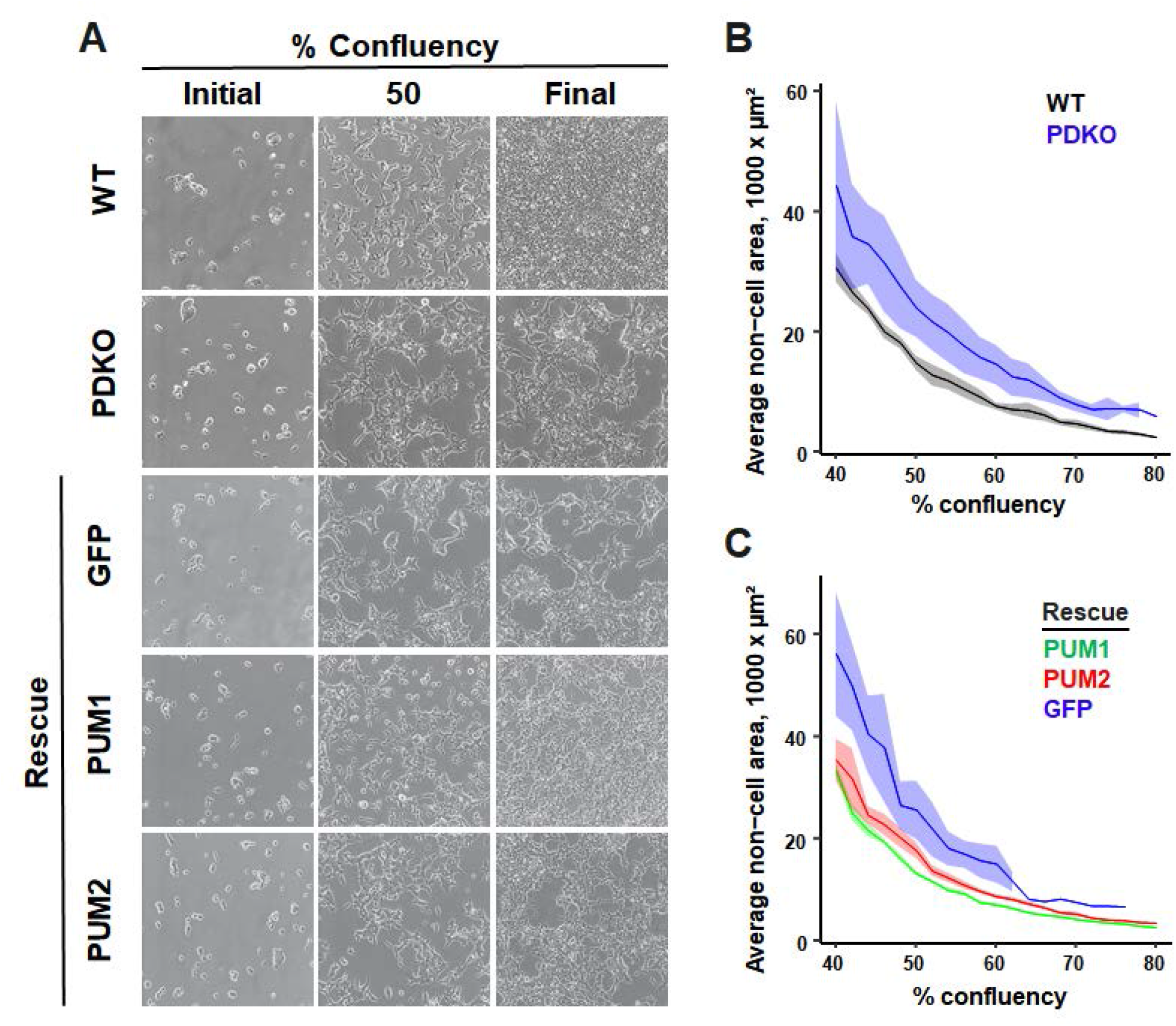
PDKO cells have cell-cell adhesion and separation defects. (A) Representative images of WT, PDKO, and rescue constructs at initial seeding, 50% confluency, and final confluency (B) Quantification of non-cell area for WT and PDKO T-REx-293 cells by % confluency. Shaded regions represent standard error of the mean. (C) Stable integrant populations of PUM1 (green), PUM2 (red), or GFP (blue) in PDKO T-REx-293 cells were analyzed for average non-cell area. Shaded regions represent standard error of the mean.

To quantify this phenotype, time-lapse images of WT and PDKO cells were collected over the course of monolayer growth, and the size of each non-cell area (individual contiguous “holes” in-between cells or groups of cells) were computed for each frame. At every timepoint, PDKO cells showed a substantially higher average non-cell area compared to WT cells (Figure 4B), reflecting their clustering phenotype. Reintroduction of either PUM1 or PUM2 abolished the clumped appearance and was accompanied by a decrease in the average non-cell area to near-WT levels (Figure 4A,C). This demonstrates that the effect on cell adhesion is PUM-specific. Differences between WT, PDKO and rescue lines were also observed in the number (as opposed to size) of non-cell areas (Supplementary Figure 3A,B), with PDKO cells showing fewer (and larger) non-cell areas. Despite the decrease in non-cell area, each rescue took longer than WT to reach maximum confluency, indicating that the growth rate defect remained (at least in part). Interestingly, the PUM1 rescue restored the cells’ ability to form a monolayer more so than PUM2, suggesting that PUM1 plays a larger role in regulating cell adhesion. GFP-expressing PDKO cells behaved nearly identically to untransfected PDKO cells, which ruled out secondary effects of transfection/selection on the measured attributes.

Time-lapse videos of WT, PDKO, and PUM1/PUM2 rescue cells were further examined to understand the clumping phenotype. During random motility in WT cultures, cells encountered other cells and made transient or prolonged contacts, but often dissociated from neighboring cells to continue independent movement. In contrast, individual PDKO (and PDKO-GFP) cells exhibited qualitatively similar motility, but typically formed stable contacts after encountering other cells, leading to aggregation into larger clusters over time (Supplementary Videos 1, 2, and 3). Most cells at the periphery of large PDKO clusters were observed to make protrusions indicative of movement away from the cluster, but remained bound, suggesting that increased cell-cell adhesion was the underlying cause of clumping. Unlike the PDKO line, PUM1 and PUM2 rescue cells were more able to detach after cell-to-cell contacts were made (Supplementary Videos 4 and 5). Taken together, these results demonstrate that PUM proteins control the inter-related properties of cell morphology, migration and adhesion, and this effect can be decoupled from PUM-dependent changes in growth.

### Extracellular matrix deposition relieves cellular aggregation

Changes in cadherin expression are often correlated with changes in adhesion between neighboring cells. To test whether an increase in E-cadherin or N-cadherin could be responsible for the increased adhesion observed, levels of both proteins were measured by western blot (Supplementary Figure 4A,B,C). E-cadherin was undetectable in both WT and PDKO cells, and there was no significant change in N-cadherin expression between cell types. Additionally, we found no differences in the levels of several proteins involved in controlling cell morphology and adhesion (Supplementary Figure 4D-I).

To assess whether the clustering behavior is related to extracellular matrix components, a thin layer of extracellular matrix (Matrigel) was deposited on the attachment substrate. After cell plating, the clumped appearance of PDKO cells reverted to the WT (Figure 5A). Additionally, culturing PDKO cells on plates where WT or PDKO cells had previously grown and deposited extracellular matrix partially relieved the clustering phenotype (data not shown). We also tested migration rates of WT and PDKO cells with and without Matrigel in the clonal ring assay. Interestingly, the addition of Matrigel had no effect on the WT migration rates and did not rescue the slower migration of PDKO cells (Figure 3E), suggesting that the PDKO decrease in bulk migration is not fully explained by the same mechanisms that are responsible for cell clustering.

**Figure 5:**
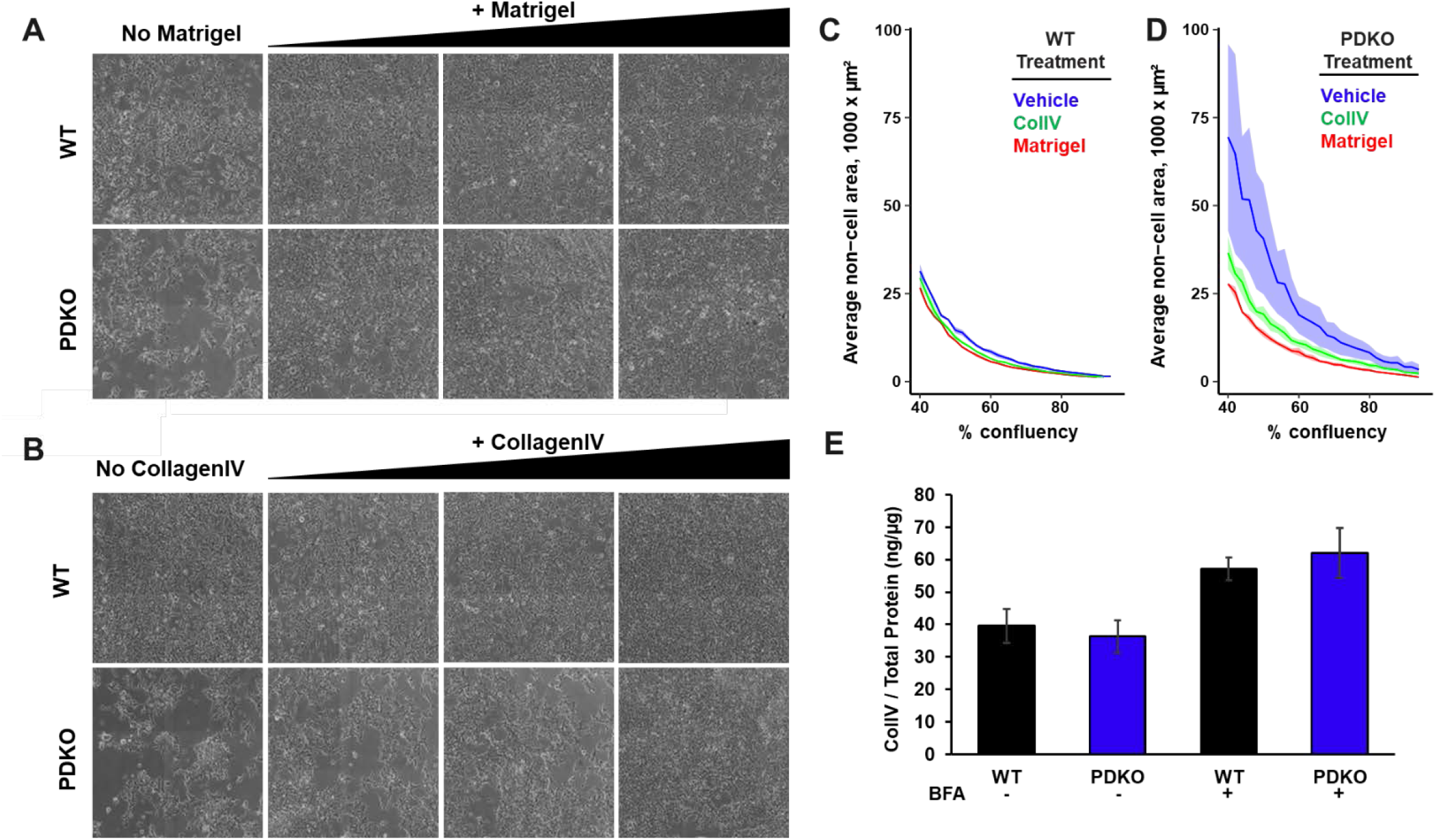
Extracellular matrix deposition relieves cellular aggregation. (A) WT and PDKO T-REx-293 cells were grown for 100 hours with varying dilutions of Matrigel, ranging from 1/100 to 1/20. (B) WT and PDKO T-REx-293 cells were grown for 100 hours with varying dilutions of ColIV, ranging from 1-10 µg per cm^2^ of growth area. (C) Quantification of data visualized in (A) and (B) for WT T-REx-293 cells grown on vehicle (blue), CollagenIV (green), and Matrigel (red). Shaded regions represent standard error of the mean. (D) Quantification of data visualized in (A) and (B) for PDKO T-REx-293 cells grown on vehicle (blue), CollagenIV (green), and Matrigel (red). Shaded regions represent standard error of the mean. (E) ELISA quantification of ColIV protein compared to total protein in WT and PDKO T-REx-293 cells with and without the addition of BFA. Measurements were calculated using 3-5 biological replicates. Error bars represent standard deviation.

To identify the molecular causes of the ECM effect, WT and PDKO cells were independently plated on two of the primary components of Matrigel, laminin and collagen type IV (ColIV). Plating on laminin exacerbated cellular aggregation in both instances (data not shown), whereas seeding on ColIV relieved the clustering phenotype for PDKO cells (Figure 5B). The rescue effect of Matrigel and ColIV was quantified using the time-lapse image metrics outlined above (Figure 5C and D). While plating WT cells on Matrigel and ColIV yielded no change in growth and morphology, plating PDKO cells on ColIV significantly relieved clumping, and Matrigel completely rescued the phenotype. Taken together, PUM-dependent effects on cell adhesion can be rescued by extracellular matrix deposition, and ColIV is a primary driving factor in this process.

The interdependence of extracellular matrix sensing and deposition with cell morphology, adhesion and migration is well known, and several scenarios may explain the observed effects. One possibility is that PDKO cells are unable to properly produce or secrete an extracellular matrix, which leads to a change in their attachment and migration properties. Another possibility is that adhesion to abundant extracellular matrix, or receptor binding to the factors within it, activates a signaling pathway that can compensate for aberrant cell-cell adhesion in PDKO cells. Given these observations, we investigated whether ColIV levels were disrupted in PDKO cells.

To separate potential effects on production from secretion, we measured ColIV levels with and without inhibition of protein transport by Brefeldin A (BFA). When measured by ELISA, we found no significant difference in ColIV protein levels between WT and PDKO cells (Figure 5E). Similar observations were made by western blot (Supplementary Figure 5A,B). Thus, the abnormal cell adhesion/motility in PDKO cells is not caused by improper ColIV production, but may be mediated by a defect in their capacity to sense ECM attachment that is compensated by additional ColIV deposition.

## Discussion

In this study, we characterized a novel role for mammalian PUM proteins in cell adhesion and migration. In addition to a previously known role in proliferation, we demonstrated an effect of PUM1+2 knockout in T-REx-293 cells on actin cytoskeleton morphology, bulk migration (clonal ring assay), and quantified a clustering/clumping phenotype that is relieved by ECM deposition. A function of PUM proteins in these inter-related processes is supported by several existing lines of evidence (reviewed in (37)). Drosophila Pum controls cell morphology at neuronal synapses (38), and mammalian Pum2 regulates dendritic outgrowth and spine morphology (39). Experiments that looked for Pum2 targets in mouse embryonic cortex by RIP (23), and for Pum1 and Pum2 targets in mouse neonatal brains by iCLIP (19), both identified an enrichment for cell adhesion and migration GO categories. Similarly, a study that used a combination of RNA-seq, CLIP/RIP-seq and bioinformatic prediction data to identify high-confidence PUM targets in HEK293 cells found an enrichment of genes involved in cell adhesion and migration under regulation (11).

Time-lapse imaging revealed that PDKO cells predominantly remained attached to each other once cell contact was made, whereas WT cells formed transient interactions and were able to detach and continue individual movement. This suggests that the identified clumping phenotype is due to defects in cell adhesion. Although we observed no differences in expression levels of E-cadherin and N-cadherin, changes in their localization to the plasma membrane and/or protein modifications could occur in PDKO cells, affecting adhesion. We have not tested all cadherins present in T-REx-293 cells, although the remaining cadherins are unconventional and are expressed at low levels in this cell type. Similarly, we have also not ruled out other proteins that contribute to or regulate cell junctions.

Additional co-culture experiments with RFP-labeled WT cells and GFP-labeled PDKO cells were also able to rescue the clumping phenotype (Supplementary Figure 6), indicating that PDKO cells in the presence of WT cells were more able to break cell-cell adhesions. Furthermore, PDKO cells did not segregate into distinct spatial areas from WT cells, which lends evidence against expression of different cadherin types. This, along with the Matrigel/ColIV experiments, suggests that misregulation of a secreted protein, or ability to interact with it, may underlie the observed phenotypes. The cycle of adhesion and de-adhesion of integrins to the ECM plays a role in actin cytoskeletal reorganization, which mediates cellular migration (40, 41). Interestingly, integrin α1β1 preferentially binds collagen IV. Thus, one possibility is that deletion of PUM1+2 affects the abundance or activity of integrin α1β1, thus leading to a clumping phenotype. It is important to note that addition of extracellular matrix could only rescue the clumping phenotype and did not rescue the bulk migration rate defect (Figure 3E), indicating that the two phenotypes are separable and are likely rooted in misregulation of different factors.

Finally, changes in actin morphology suggest that PUM proteins may impact the activity of the Rho family of GTPases. These pathways, along with controlling actin structure, are known to regulate cell motility and cell adhesion (42). Thus, GTPase activating proteins and guanine exchange factors, along with the GTPases themselves, are potential PUM targets, and an enrichment of related gene ontology categories among PUM targets has been reported (11, 19).

Overall, the processes regulating cytoskeletal morphology, cell-cell adhesion and interactions with the ECM are intimately interdependent through multiple feedback loops. It is therefore not surprising that PUM proteins affect these processes together, and the detailed molecular mechanisms that link them to PUM regulation await further investigation. Manual examination of transcripts that are differentially expressed in PUM DKO cells, partially restored upon rescue (this study), bound by PUM in CLIP-seq experiments (10, 43), and contain the PUM binding motif in their 3‵ UTRs, identified two candidates that are associated with these processes. Frizzled-8 (FZD8) is a non-canonical Wnt protein receptor (44), and this signaling pathway is known to play a role in the regulation of cell adhesion and migration (45). NECTIN-4 belongs to a class of proteins involved in cell-cell adhesion, and expression of NECTIN-4 on the surface of ovarian cancer cells increases adhesion, although its expression is also associated with increased migration in a scratch assay (46).

In our experiments, PUM1 and PUM2 showed partially overlapping, but non-identical roles in regulating growth and cell migration/adhesion, in agreement with complementary roles in embryogenesis (47). Each protein was able to partially rescue the observed defects, indicating some functional overlap. On the other hand, since neither PUM protein was sufficient for compete rescue, some specific functions likely remain. This idea is supported by the fact that distinct sets of bound targets have been previously found (10, 12, 19, 47) Alternatively, this result may be explained by dosage effects(30), since total PUM expression levels in individual rescues are less than in WT cells. Additionally, we note that PUM1 displays a somewhat increased ability to rescue growth and adhesion phenotypes, compared to PUM2. This result is also reflected in the RNA-seq data, where more genes were differentially expressed upon rescue with PUM1 that PUM2. Since western blots confirm that both PUM1 and PUM2 were rescued to approximately WT levels (Figure 1A), this effect is likely specific to PUM1. Interestingly, PUM1 is routinely found to interact with a larger set of mRNAs than PUM2 (10, 12, 19, 47).

Adhesion and migration play important roles in coordinating many cell behaviors and processes. Proper control of cell-cell adhesion and migration are essential for cell organization during development, and cancer cells are known to acquire malignancy through decreased contacts with neighboring cells and upregulating motility genes (48, 49). Understanding the role of PUM proteins in in these processes is essential for providing better models for its function in developmental processes, as well as understanding how changes in PUM expression can influence cancer.

## Materials and Methods

### Cell culture

T-REx-293 cells were obtained from Invitrogen. HCT116 cells were a gift from the Mendell lab(30). PUM double knockout cells were generated as described previously (10). T-REx-293 cells were grown in DMEM (Corning) with 10% fetal bovine serum (Corning) and 10 units/mL of penicillin/streptomycin (Gibco) at 37 °C with 5% CO2. HCT116 cells were grown in McCoy’s 5A (Iwakata & Grace Modification) media (Corning) with 10% fetal bovine serum (Corning) and 10 units/mL of penicillin/streptomycin (Gibco) at 37 °C with 5% CO2. For the addition of Matrigel Matrix (Corning), 1 mL total volume of ice-cold diluted Matrigel was added to a 6-well and incubated for 1 hour at room temperature. Excess solution was removed just prior to plating. Appropriate ice-cold media was used to dilute Matrigel from 1/1000 to 1/20. Purified mouse collagen IV (Corning) was diluted in ice-cold sterile DI water containing 0.05 M HCl, added to a 24-well plate, and incubated at room temperature for 1 hour. Wells were aspirated and washed with 500 µl PBS for 5 minutes at room temperature. Plating dilutions used were between 1-30 µg/cm^2^ growth area. Protein transport inhibition in T-REx-293 cell culture was achieved by the application of 1 µl/ml BD GolgiPlug (Fisher Scientific), diluted in appropriate culture medium. An equivalent amount of DMSO was used as a vehicle control. Cells were incubated for 5 hours before being trypsinized and washed 3 times with ice-cold PBS. Centrifugation was performed at 1500 xg for 5 minutes. Cell pellets were then used in downstream western blot and ELISA analysis.

### Rescue cell lines

Overexpression plasmids for PUM1 (pLX302-PUM1), PUM2(pLX302-PUM2), and GFP (pLX302-GFP) were a gift from the Mendell lab (30). TransIT-LT1 reagent (Mirus) was used per manufacturer’s instruction to add 1 µg of plasmid to PUM double knockout T-REx-293 cells seeded in 6-well plates at ∼70% confluency. 48 hours after transfection, cells were selected with 1 µg/mL puromycin for at least 7 days. PUM1 and PUM2 expression was confirmed by western blot using a goat anti-PUM1 antibody (Bethyl, A300-201A) and a rabbit anti-PUM2 (Bethyl, A300-202A).

### Cell growth measurements

Cells from an 80-90% confluent plate were trypsinized, counted on a hemocytometer and plated at an initial density of 325,000 cells in multiple wells of 6-well plates. At regular intervals, individual wells were harvested for counting on a hemocytometer, averaging 2-3 1 mm x 1 mm hemocytometer squares for each biological replicate. A total of 3 biological replicates were measured. Growth parameters were derived from standard logistic growth nls model fits in R. Statistical significance of differences in model fits between cell types were determined by pairwise F-test ANOVA comparisons of nested models that incorporate or ignore the cell type, as described (50).

### Image analysis of time-lapse microscopy images

For time-lapse microscopy of WT, PDKO and rescue lines (Figure 4), cells from an 80-90% confluent plate were trypsinized, counted on a hemocytometer and plated at an initial density of 325,000 cells in a 6-well plate 24 hours prior to imaging. Phase contrast images were collected with a Biostation CT over 72 hours every 10-15 minutes. A total of 6 biological replicates for each cell type, consisting of 2-3 separately imaged regions (“frames”, technical replicates) per each cell type / biological replicate were collected. For time-lapse microscopy of Matrigel or collagen-treated samples (Figure 5), cells were trypsinized and counted as above, then plated at an initial density of 30,000 cells per well in a 24 well plate 24 hours prior to imaging (ColIV and Matrigel treatment is described in “cell culture”). Phase contrast images were collected with a Biostation CT every hour for 120 hours. A total of 3 biological replicates for each cell type were collected. Each biological replicate consisted of 2 technical replicates. All wells were imaged in a 4×4 tile using a 10x objective.

Individual areas occupied by cells and by empty space between cells, as well as overall confluency, were computed by the available functions in CL Quant software (Nikon). Frames that started growing at >50% confluency and that never reached >55% confluency were filtered out of the analysis, as they behaved differently over the time course. Imaged frames were then aligned by confluency, and biological replicates were averaged together using R.

For the calculation of the number of non-cell areas as a function of time (as opposed to confluency), the time axes of image series were aligned to each other. Individual biological and technical replicates exhibited trajectories of computed metrics over time that were very stereotypical within a given cell type, but were shifted relative to each other along the x (time) axis. The shifts resulted from stochastic variability in local seeded density between the imaged frames, and a variable cell recovery lag phase after the seeding of cells and prior to the onset of active growth and motility. To eliminate this variability, the time axes of individual frames within a given cell type were shifted such that the cell areas (confluency) in the frames were aligned to each other with maximal overlap using the dtw R package with a rigid step pattern and averaged within each cell type. This operation thus re-aligned the time axis of each technical replicate based on having the same growth stage (cell area or confluency). To compare between cell types, the time axes of averaged traces were aligned to each other over the initial 10% of their respective growth curves, to be able to analyze subsequent growth from a starting point of equivalent cell area.

### Clonal ring migration assays

Cells from an 80-90% confluent plate were trypsinized, counted on a hemocytometer and plated at an initial density of 800,000 or 1,000,000 for T-REx-293 and HCT116 cells, respectively, inside a 6 mm I.D. cloning ring within a 6-well. Cells were given 4-6 hours to attach to the plate before the cloning ring was removed. Images were taken every 12 hours for 60 hours to track collective migration of cells. Images were processed, aligned, and measured in ImageJ. For migration assays supplemented with Matrigel, a 1/50 dilution was used (protocol as described above).

### Fluorescence Microscopy

TransIT-LT1 reagent (Mirus) was used per manufacturer’s instruction to add 1 µg of the mRFP-UtrCH plasmid (Addgene #26739) to both WT and PDKO T-REx-293 cells seeded in 6-well plates at ∼70% confluency. Cells were trypsinized and resuspended during transfection in order to increase transfection efficiency. 48 hours after transfection, cells were trypsinized and plated onto NaOH (2 M for 2 hours) and poly-lysine (0.5 mg/mL on shaker for 1 hour) treated glass-bottom plates. Live cells were imaged 24 hours post seeding using a custom-built spinning disk confocal microscope (Solamere Technology) with a Yokagawa W1 spinning disk (Yokagawa), EM-CCD camera (Hamamatsu 9100c), and a Nikon Eclipse TE (Nikon) inverted stand. A 60× water immersion lens (1.2 NA) was used with perfluorcarbon immersion liquid (RIAAA-678, Cargille). The stage is fully motorized and controlled by Micromanager software (www.micromanager.org) with ASI Peizo (300-μm range) and a 3 axis DC servo motor controller. Solid-state lasers (Obis from 40 to 100 mW) and standard emission filters (Chroma Technology) were used. A 561 laser with emission filter 620/60 was used.

### RNA-seq library preparation and analysis

For each cell type, three biological replicates were collected and processed separately. Cells were cultured to 50% confluency in a 10 cm plate. Total RNA was extracted with Ribozol, and libraries were prepared using the NEB mRNA magnetic isolation module (E7490S), NEBNext Ultra RNA Library prep kit (E7420L), and NEB Multiplex Oligos for Illumina Index Primer Sets 1 (E7335S) and 2 (E75500S) and sequenced on an Illumina NextSeq instrument. Further processing was done in R: reads were aligned to the GRCh38 genome assembly using HISAT2 and annotated with Gencode v40 annotations. Differentially expressed genes were identified using DESeq2. The sequencing data is available at GEO series record GSE207836.

### Western blots

For collagen IV western blots, primary rabbit anti-Collagen IV (abcam ab6586) and secondary anti-rabbit IgG, HRP linked (Cell signaling technology) antibodies were used. T-REx-293 cells were plated at 30% confluency in 10 cm plates and allowed 24 hours to adhere. Cells were incubated with BD GolgiPlug (Fisher Scientific) or DMSO for 5 hours and collected at ∼60% confluency and run on an 8% SDS-polyacrylamide gel. Gels were wet transferred, overnight at 4 °C, to a PVDF membrane. The following day all steps were done shaking at RT, membranes were blocked in 5% powdered non-fat milk TBST for 1 hour, then incubated with primary antibody for 1 hour, washed in TBST for 5 minutes 3 times, incubated with secondary antibody for 1 hour, and washed in TBST for 5 minutes 3 times. Finally, membranes were visualized by Radiance Q (Azure Biosystems AC2101). Blots were imaged using a ChemiDoc imaging system.

Antibodies used for candidate western blots are as follows: mouse anti-E-cadherin (BD Transduction Laboratories, 610181 and Santa Cruz, sc-8426), mouse anti-N-cadherin (Santa Cruz, sc-393933), mouse anti-ephrin-B1 (Santa Cruz, sc-515264), mouse anti-Wnt-5a (Santa Cruz, sc-365370), mouse anti-c-Jun (Santa Cruz, sc-74543), mouse anti-GSK-3a (Santa Cruz, sc-5264), and mouse anti-BPIX (Santa Cruz, sc-393184). T-REx-293 and HCT116 cells were collected at ∼70% confluency and run on an SDS-polyacrylamide gel. Gels were transferred to a nitrocellulose membrane, blocked, and incubated with primary antibody overnight at 4 °C.

### ColIV ELISA

ColIV protein quantification by ELISA was conducted using a COL4 ELISA kit (MBS2701454, MyBioSource) per manufacturer’s specifications. In brief, T-REx-293 cells were collected after BD GolgiPlug treatment, as described above, and lysed using a standard 1% NP-40 buffer (150 mM NaCl, 1% total volume NP-40, 50 mM Tris-Cl pH 8.0). Lysate was centrifuged at 1500 xg for 10 minutes at 4 °C, aliquoted and stored until needed at -80 °C. Lysates were limited to one freeze-thaw. 100 µl of undiluted lysate was used for all samples. ELISA samples were read on a Tecan Spark spectrophotometer at 450nm. Total protein was retroactively calculated using the DC protein assay (Bio-Rad) according to the manufacturer’s specifications. DC Protein assays were read on Tecan Spark at 750nm.

### Mixing Experiments

The pCMV DsRed-Express2 (Clontech 632539) expression plasmid was stably transfected into wildtype T-REx-293 cells by same protocol described above and was selected using Neomycin at 500 µg/mL for seven days. The pMSCV-PIG (Addgene #21654) expression plasmid was stably transfected into wildtype 293T cells by the same protocol described above and was selected using Puromycin at 1 µg/mL for seven days. Generation of T-REx-293 PDKO+GFP cells is described above. Cells were co-cultured at a ratio of 1 to 1 and a total cell volume of 325,000 cells per 6-well. Time-lapse images were collected as described above.

## Acknowledgements

We thank Prue Talbot, Rachel Behar and the UCR Stem Cell Center for time-lapse imaging assistance, and the Rasmussen lab for microscopy help. We thank the Riccomagno lab for utrophin constructs. This work was supported in part by NIH grant 1R21NS118390-01.

## Supplementary Figures

**Supplementary Figure 1:**
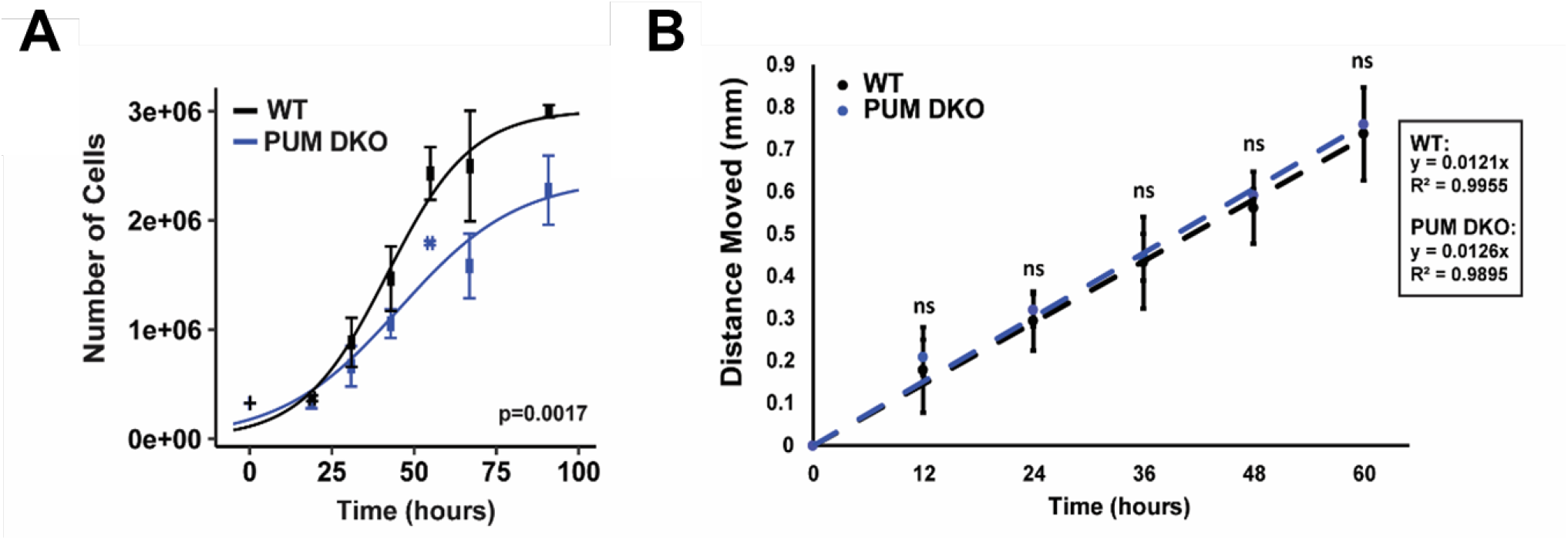
HCT116 PUM DKO cells show growth defects, but not strong adhesion and migration defects. (A) Growth rates of WT (black) and PUM DKO (blue) HCT116 cells. Growth measurements were calculated from three biological replicates, with error bars representing standard error of the mean. For the purposes of plotting, the time points are grouped for each cell type / replicate group (since the data was collected at slightly different timepoints for each replicate), generating error bars along the x axis direction (standard error of the mean). Logistic growth model fits of the data were compared using ANOVA F-test to determine statistical significance. (B) Migration rates of WT (black) and PUM DKO (blue) cells. Measurements were calculated over 3 biological replicates. Statistical significance was determined by Student’s T-test.

**Supplementary Figure 2:**
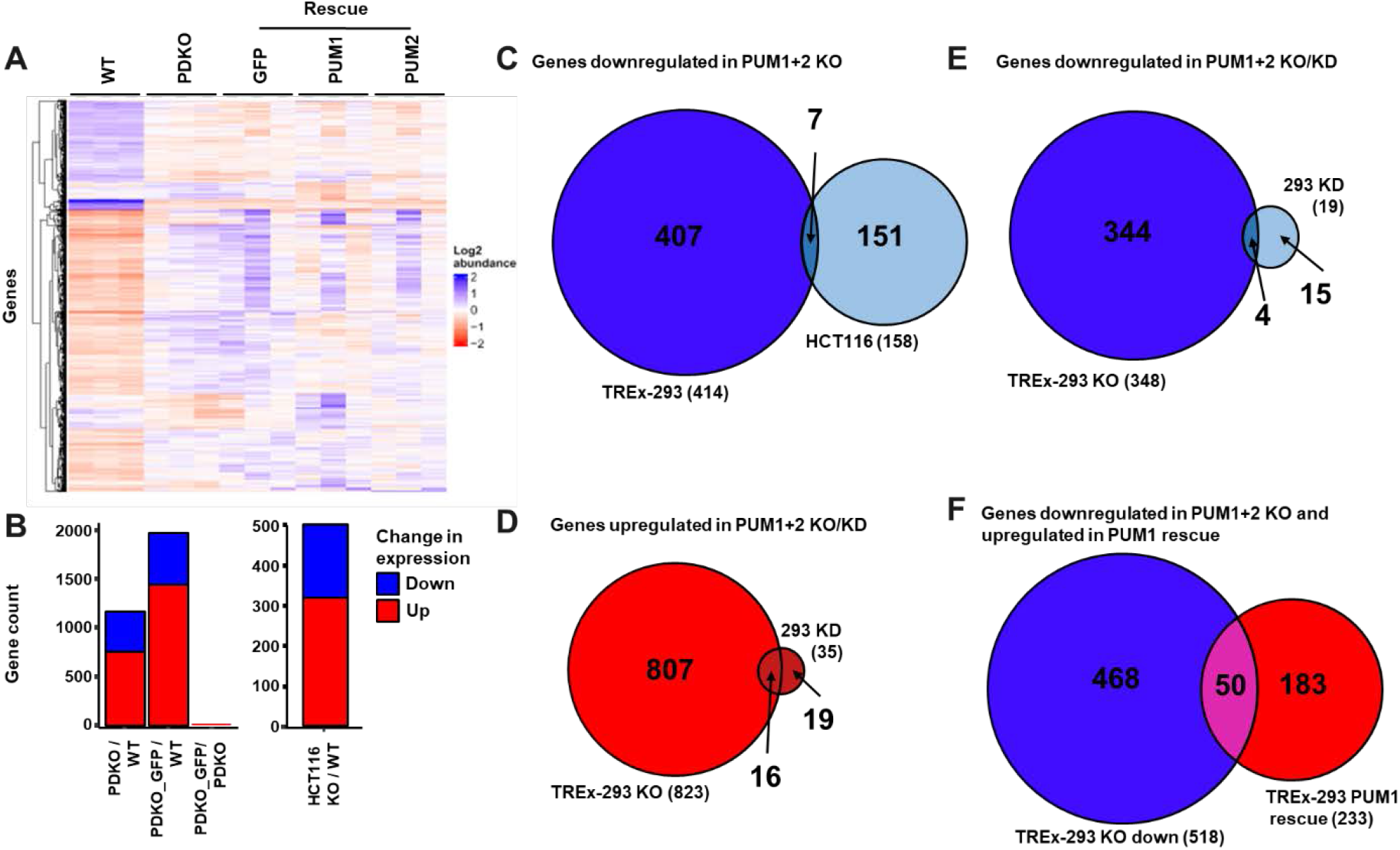
(A) Heatmap of RNAseq gene expression changes between WT, PDKO and rescue T-REx-293 cells. (B) Number of differentially expressed genes in T-REx-293 PDKO vs WT, PDKO+GFP vs WT, and PDKO+GFP vs PDKO cells (left), and in HCT116 PDKO vs WT cells (right). (C) Overlap among genes downregulated in PDKO vs WT T-REx-293 and HCT116 cells. (D) Overlap among genes upregulated in PDKO vs WT T-REx-293 cells and 293 PUM1+2 KD cells. (E) Overlap among genes downregulated in PDKO vs WT T-REx-293 cells and 293 PUM1+2 KD cells. (F) Overlap of genes downregulated in T-REx-293 PDKO vs WT with genes upregulated in PUM1 rescue vs PDKO.

**Supplementary Figure 3:**
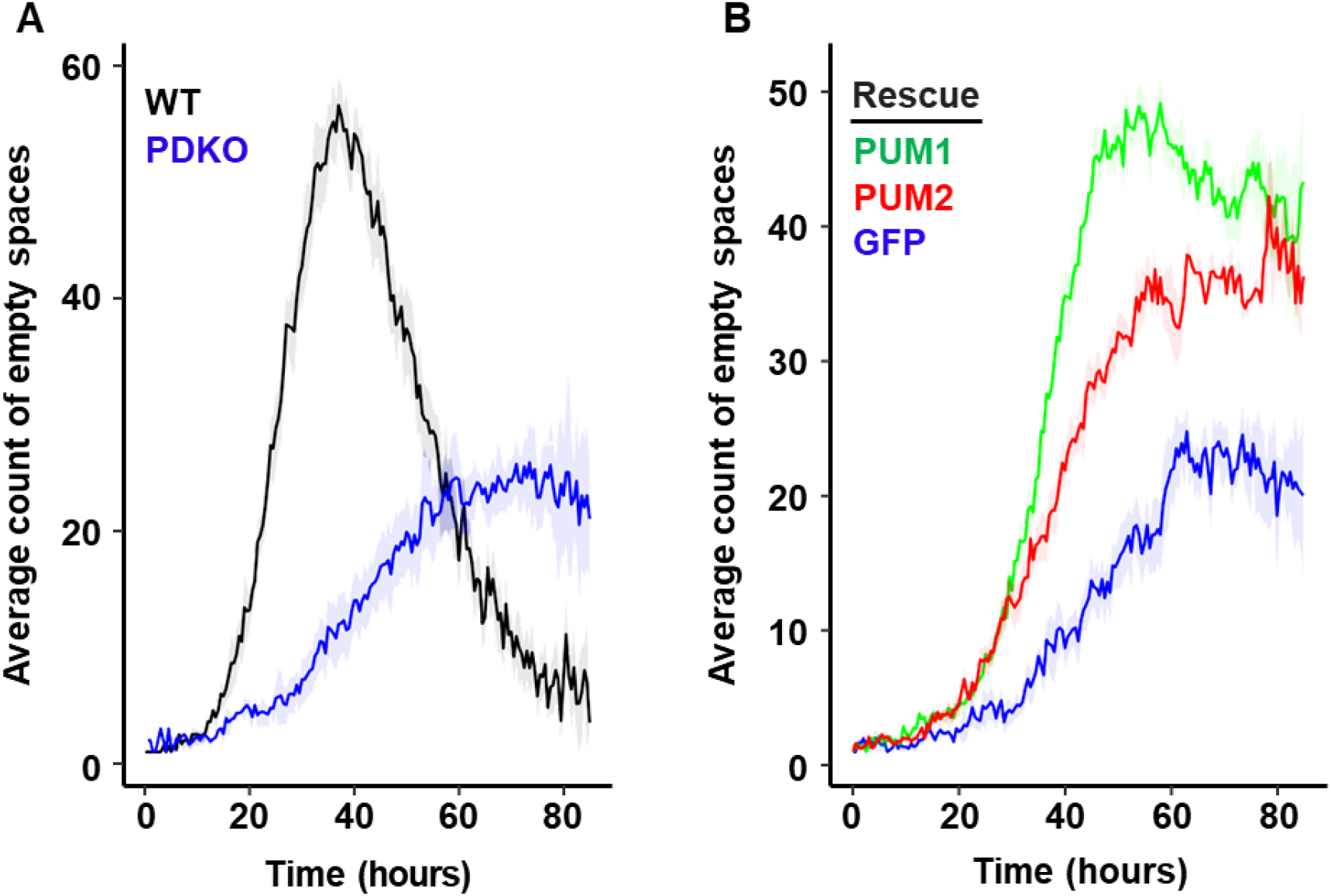
PDKO T-REx-293 cells are unable to form a monolayer typical of WT T-REx-293 cells and this phenotype can be alleviated by the exogenous expression of PUM1 or PUM2. (A) WT (black) and PDKO (blue) T-REx-293 average number of space objects. (B). Stable integrant populations of PUM1 (green), PUM2 (red), or GFP (blue) in PDKO T-REx-293 cells were analyzed for average number of space objects. Measurements were calculated from six biological replicates. Shaded band represents two standard deviations from the mean.

**Supplementary Figure 4:**
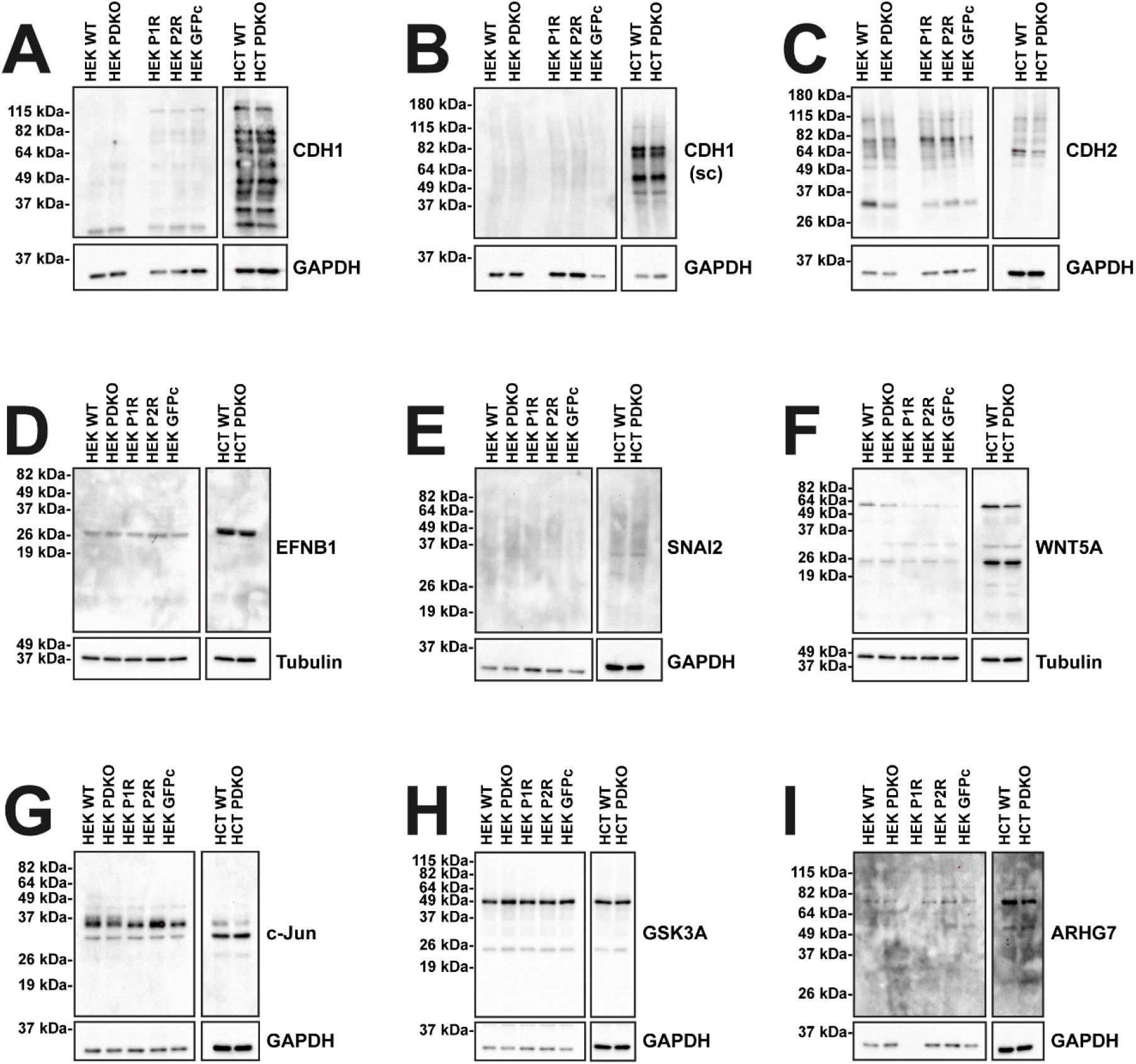
The PDKO adhesion phenotype is not caused by changes in cadherin expression levels, or levels of select candidate genes. Western blots across WT, PDKO, and rescue T-REx-293 cells for (A and B) E-cadherin, (C) N-cadherin, (D) Ephrin B1, (E) Snail2, (F) Wnt5A, (G) c-Jun, (H) GSK-3α, and (I) Arhgef7.

**Supplementary Figure 5:**
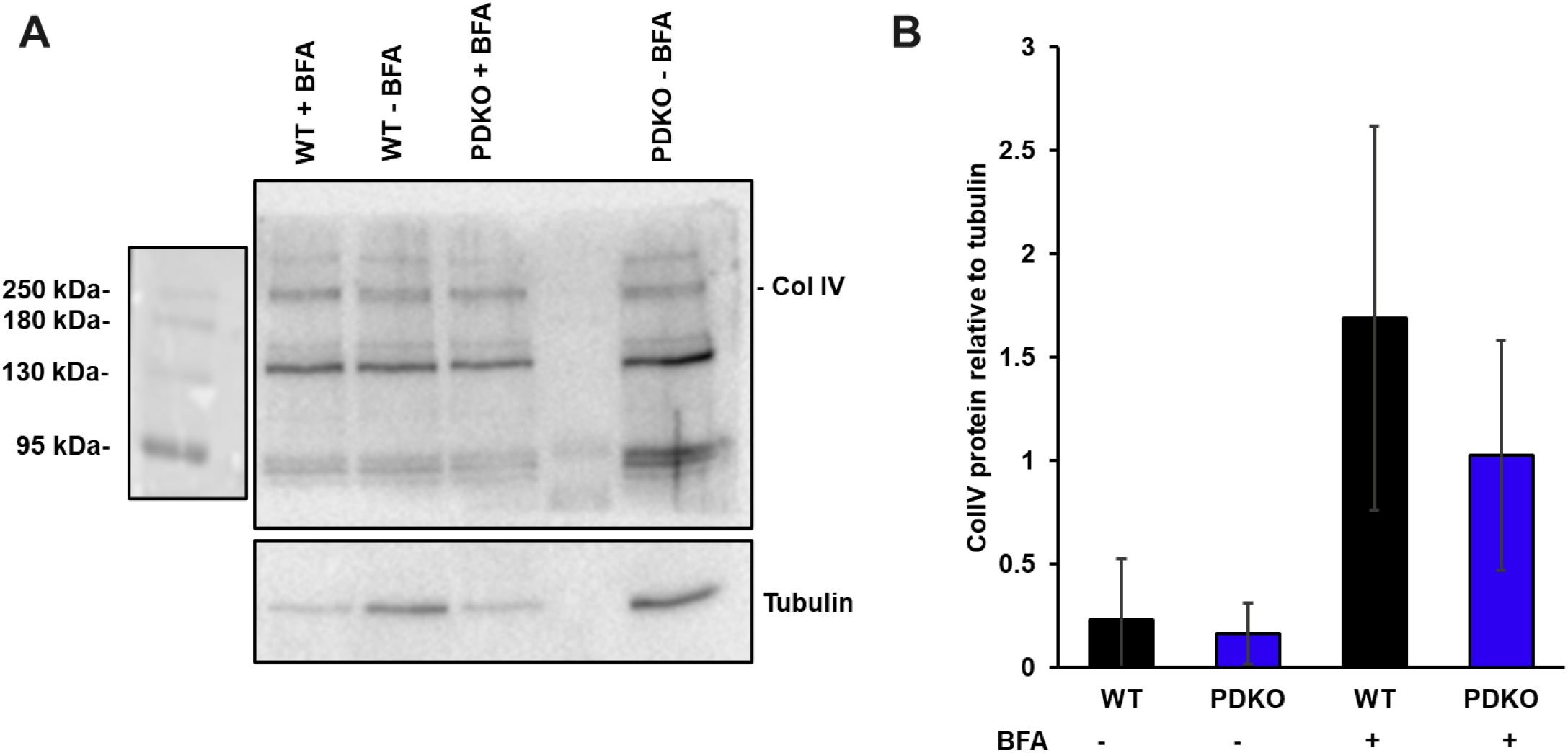
ColIV protein levels do not change upon PDKO. (A) Representative western blot image of ColIV. Tubulin serves as a loading control. (B) Western blot quantification of ColIV protein compared to total protein in WT and PDKO T-REx-293 cells with and without the addition of BFA. Error bars represent standard deviation.

**Supplementary Figure 6:**
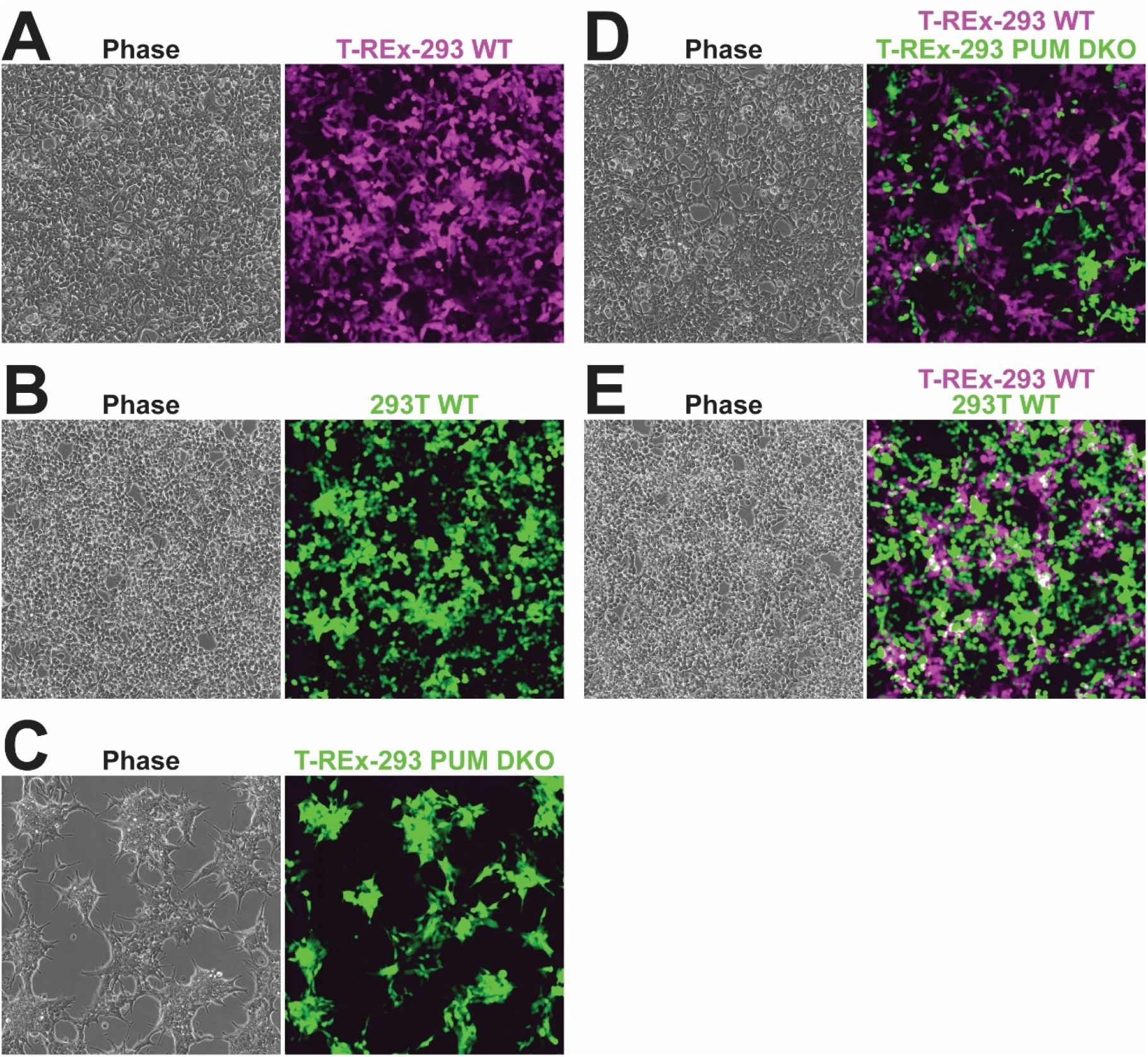
Co-culture of WT and PDKO cells rescues the increased cell adhesion phenotypes. (A) Phase contrast and fluorescence images of T-REx-293 WT only, (B) 293T WT only, (C) T-REx-293 PUM DKO only, (D) co-cultured T-REx-293 WT and PUM DKO, and (E) co-cultured T-REx-293 WT and 293T WT cells. Images were collected after 72 hours of culture.

## Supplementary Video Legends

**Supplementary Video 1: Time-lapse video of WT T-REx-293 cells**. Frames correspond to 15-minute intervals over a 72-hour period.

**Supplementary Video 2: Time-lapse video of PUM DKO T-REx-293 cells**. Frames correspond to 15-minute intervals over a 72-hour period.

**Supplementary Video 3: Time-lapse video of PUM DKO T-REx-293 cells stably expressing GFP**. Frames correspond to 15-minute intervals over a 72-hour period.

**Supplementary Video 4: Time-lapse video of PUM DKO T-REx-293 cells stably expressing PUM1**. Frames correspond to 15-minute intervals over a 72-hour period.

**Supplementary Video 5: Time-lapse video of PUM DKO T-REx-293 cells stably expressing PUM2**. Frames correspond to 15-minute intervals over a 72-hour period.

